# Arousal levels explain inter-subject variability of neuromodulation effects

**DOI:** 10.1101/2020.05.08.083717

**Authors:** Marco Esposito, Clarissa Ferrari, Claudia Fracassi, Carlo Miniussi, Debora Brignani

## Abstract

Over the past two decades, the postulated modulatory effects of transcranial direct current stimulation (tDCS) on the human brain have been extensively investigated, with attractive real-world applications. However, recent concerns on reliability of tDCS effects have been raised, principally due to reduced replicability and to the great interindividual variability in response to tDCS. These inconsistencies are likely due to the interplay between the level of induced cortical excitability and unaccounted individual state-dependent factors. On these grounds, we aimed to verify whether the behavioural effects induced by a common prefrontal tDCS montage were dependent on the participants’ arousal levels. Pupillary dynamics were recorded during an auditory oddball task while applying either a sham or real tDCS. The tDCS effects on reaction times and pupil dilation were evaluated as a function of subjective and physiological arousal predictors. Both predictors significantly explained performance during real tDCS, namely reaction times improved only with moderate arousal levels; likewise, pupil dilation was affected according to the ongoing levels of arousal. These findings highlight the critical role of arousal in shaping the neuromodulatory outcome, and thus encourage a more careful interpretation of null or negative results.

## 1. Introduction

Founded on decades of experimentation, transcranial direct current stimulation (tDCS) is a research tool capable of interacting with the central nervous system, that has been rediscovered at the beginning of this century (1). Beside its value for basic research (2), tDCS has raised great interest for real-world applications, like rehabilitative interventions for neurological and psychiatric diseases (3) and cognitive enhancement (or detraction) in both young and older adults (4–7). However, the development of more effective and generalizable stimulation protocols has been hindered by the gap between our sparse knowledge of the physiological effects and the induced behavioral impact of tDCS (8). What raises most concern is the lack of replicability among tDCS studies and the interindividual variability in response to tDCS (9–15). In addition to non-optimal methodological practices, such as inadequate control conditions and lack of statistical rigor, a complex interplay among biological differences and the level of neuromodulatory effects might be crucial in explaining the reported inconsistencies across studies (16–18). In particular, state-based factors, including the specific or generalized levels of activation prior and during stimulation, the initial levels of performance, wakefulness, task priming or novelty, might all play a decisive role. It appears conceivable to interpret the final effects of tDCS as contingent on the level of network engagement (19,20). In line with this prediction, several cognitive studies have reported a clear effect of baseline levels of different mental capabilities on tDCS response (21–26). Most recently, individual differences in the behavioral effects of prefrontal tDCS have been associated with the levels of excitability of the targeted cortex, indexed by relative concentrations of GABA and glutamate (27).

Notably, tDCS affects large-scale brain systems extending well beyond the area under the stimulating electrode (28–31). This approach translates into a lack of focality that closely resembles the spread of the noradrenergic modulatory action exerted by the locus coeruleus (LC), which subtends arousal functions. Several authors have highlighted the key adaptive role of this specific midbrain system in shaping behavioral performance of primates (32–36). A large body of evidence suggests that the exogenous direct currents and the endogenous noradrenergic modulatory action on target cells, share the same central mechanism of neuronal gain control (34,37–39). Therefore, an interrelation between the two stimulating activities seems reasonable to the extent that whenever the contrast between activated and inhibited units becomes sufficiently increased or decreased any further added neuromodulation can likely spoil the expected results. In this regard, a recent study has shown that offline anodal tDCS may hinder the LC endogenous action during response inhibition processes due to the induced alterations of pre-existent neural excitability levels (40). Given the above considerations, it appears evident that great part of the tDCS behavioral variability reasonably stems from the interdependency between the induced cortical excitability and the varying levels of arousal experienced by participants before and during the experimental sessions.

The aim of this study was to verify whether the behavioural and physiological responses induced by a common prefrontal tDCS montage were dependent on the participants’ arousal levels. We selected the tDCS montage used to stimulate prefrontal cortex in attentional and vigilance tasks (40–43), which is also commonly used in a variety of other settings, such as language-related, executive functions, episodic and visual working memory tasks (44–47). The tDCS was applied during an auditory oddball task aimed to probe cognitive performance as a function of arousal levels (48–50). Our task, indeed, was purposefully designed to keep participants alerted over uncertain intervals (i.e., variable inter stimulus interval) in a way that online tDCS effects would be necessarily subjected to more frequent fluctuations of arousal (51,52).

We tracked pupillary changes as a proxy for the LC modulatory action (50,53–55). Accordingly, we used reaction times (RT) and pupil dilation peaks (PD) as measures of LC phasic response to the relevant stimuli (target), and pre-stimulus pupil diameter (PrePD) as a physiological marker of the LC tonic discharge activity. Furthermore, because LC endogenous activity is closely related to the perceived anxiety (56–58) subjective arousal levels were evaluated by means of State-Trait Anxiety Inventory (STAI-Y) (59).

## 2. Methods

Mindful that the mere sensory stimulation could mimic the expected arousal effects, prior to conducting the study, we ran a control experiment to validate our blind-controlled tDCS protocol with respect to the potential alteration of arousal due to subjective sensations. To this end, ten healthy participants were recruited. Pupillary dynamics were recorded at rest using the exact same setting as in our main experiment (see section 2.3 and 2.4). Statistical analyses revealed no difference in eye-blink rate and subjective discomfort between sham and real stimulation, ruling out the possibility of tDCS confounding effects on arousal (see supplementary material).

### 2.1 Participants

Fifteen right-handed healthy participants took part in the main experiment. Data of one subject were rejected prior to analyses due to the excessive noise in her/his pupil signal (i.e., interpolation rate > 30% of the whole epoch; see 2.3). The remaining 14 participants (8 females) had a mean age of 22.4 (*SD* = 3.9) and a mean score to the STAI-Y trait of 44.9 (*SD* = 4.1). Participants had no history of neurological or psychiatric illness and had normal or corrected-to-normal visual acuity. Ethical approval was obtained by the Ethics Committee of the IRCCS Centro San Giovanni di Dio Fatebenefratelli, Brescia, Italy. All participants were given written informed consent.

### 2.2 Study design and task procedure

A single-blind within-subject design was implemented for this experiment. The testing sessions were organized in two days separated by at least 48h in order to exclude any tDCS carryover effects. In each session we collected behavioral and pupil data for the whole task duration (~18 min). Participants completed the task twice: at *baseline* (T1) without any electrodes mounted on their scalp, and subsequently either during *sham* or *real* stimulation (T2) (Figure 1b).

**Fig. 1.**
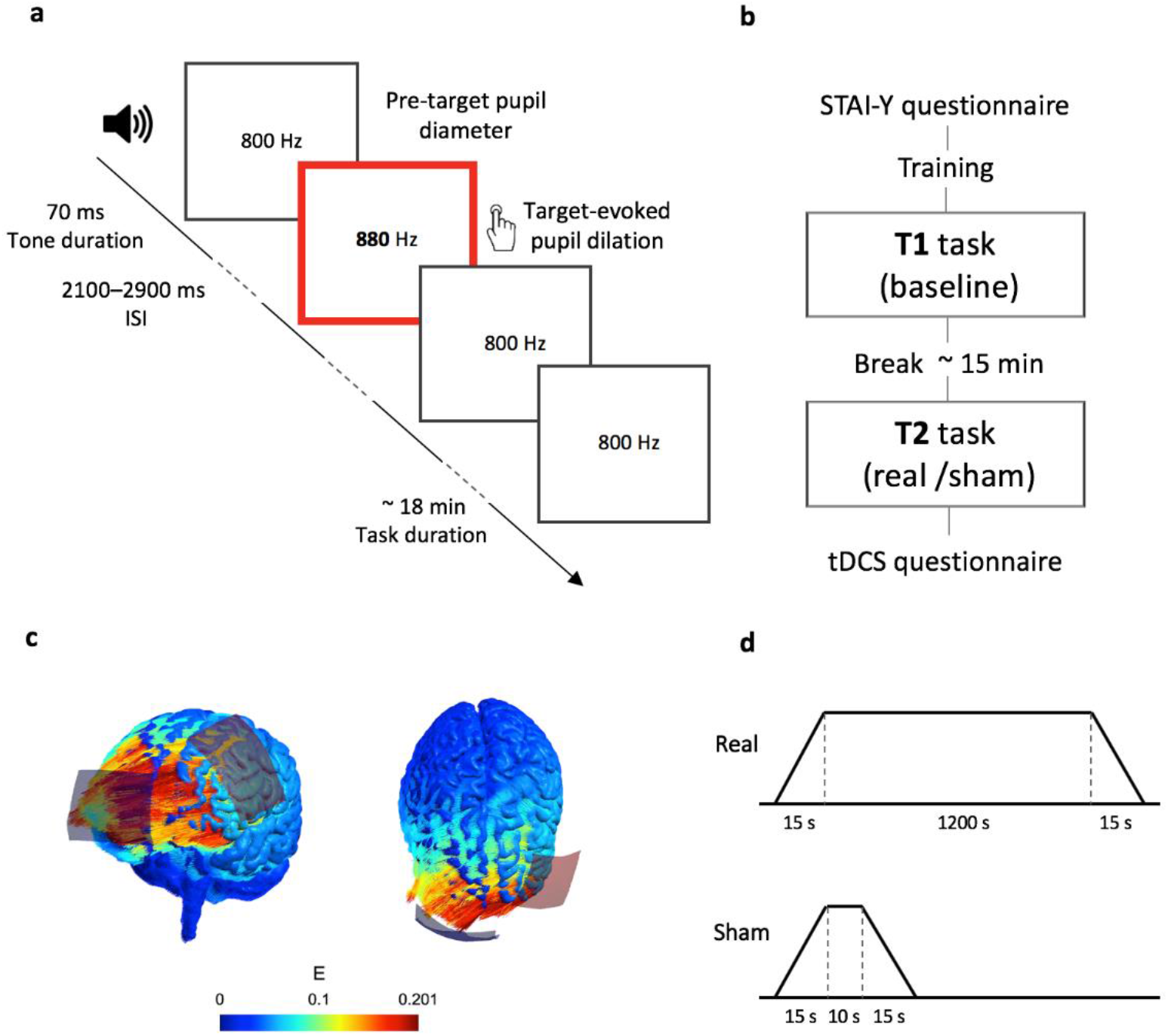
Study design and task paradigm. **a**, Example of a trial sequence. **b**, Overview of the experimental timeline, showing two testing sessions each one with two task conditions: baseline and stimulation. **c**, Simulation results for the applied tDCS montage and parameters using SimNIBS toolbox (Saturnino et al., 2019). The colors denote the electric fields simulated in a default head model. **d**, Schematic representation of the stimulation protocol

Participants were randomly assigned and counterbalanced across two session-orders of tDCS protocol, so as to rule out any extra confounding variable. Importantly, they were kept blind to the ongoing experimental condition (i.e., sham or real). However, any proportion of variability possibly due to either orders of stimulation was accounted for by including the order group as an independent fixed factor (see section 2.5). As for the time of the day, the same participant was tested at around the same hour to control for any arousal variation due to the daily metabolic cycle and circadian rhythms (60).

Participants seated in a soundproof dark room at the distance of about 55 cm from a 17-in LCD monitor and with the only source of light provided by a grey fixation cross. The auditory oddball task was presented using E-Prime presentation software (61) by means of two constant-loudness speakers (Figure 1a).

In every task condition there was a fixed total number of trials (420) of which 20% included targets (84) and 80% standards stimuli (336). The stimuli order was then pseudorandomized in a way that target tones (880Hz) occurred after at least three standard tones (800 Hz). The interstimulus interval was set to a range of 2.1-2.9 s and both stimuli lasted for 70 ms including 5 ms of fade in-out edit. In so doing, we ensured enough time (~8 s) for any pupil dilation to return to baseline before overlapping to the next target trial (50,53). Along with a short training session, participants were instructed to readily press a button with their right index finger whenever detecting a target tone, and to keep their gaze on the fixation cross throughout the task. Speed of response and gaze fixation were emphasized before each task execution.

At the end of each experimental session participants were given a questionnaire to rate the perceived sensations or discomforts that influenced their performance (62,63).

Finally, a careful screening on the amount of sleep, caffeine intake, nicotine and alcohol consumption was carried out next to the above questionnaire. None of these factors was found to be associated with either stimulation sessions.

### 2.3 Pupil signal recording and pre-processing

Participants seated on a chair with adjustable height allowing for the use of a fixed chinrest, and thus keeping variability in the eye-to-camera distance and visual angle as low as possible. For the pupil diameter recording, an EyeLink 1000 Plus system (SR Research, Osgood, ON, Canada) was set up at 500 Hz sampling rate with left-monocular and pupil-CR tracking mode. A 9-point calibration procedure was performed before each recording session. After the final session, participants were asked to wear hand-crafted goggles whose left side incorporated an artificial eye with a 4 mm pupil, carefully positioned over the subject’s left eye. This allowed for a precise conversion of pupil arbitrary units from the eye-tracker system output to millimeters. Pupil signal was processed offline. Eye blink correction was implemented with a custom script in MATLAB (MathWorks, Inc, Natick, MA, USA). A shape-preserving piecewise cubic interpolation method was chosen to interpolate values ranging from 70 ms before blink onset to 300 after blink offset. Epoch segmentation (−1 s to +2.5 s, relative to target onset), baseline correction (subtractive method, from −800 ms to +200 ms) and visual inspection of pupil traces was carried out in the Brain Vision EEG analyzer software (Brain Products GmbH, Munich, Germany). We extracted two variables of interest from pupil signal: (i) pupil dilation (PD), as the peak value of the maximum dilation after targets presentation and (ii) Pre-stimulus pupil diameter (PrePD) as the mean of 1 s data prior to tone presentation. All epochs with a peak pupil diameter exceeding ±2 mm were rejected (50).

### 2.4 tDCS protocol

A battery-driven current stimulator (Brain-STIM, EMS, Bologna, Italy) was used to deliver 1 mA (0.028 mA/cm^2^) direct current stimulation via two rubber electrodes (35 cm^2^) which were inserted inside two saline-soaked sponges. These were fixated with an elastic mesh stretching over the entire head. In order to ensure a stable impedance level as well as keeping skin sensations at the minimum, conductive electro-gel was also applied.

Similarly to previous studies (43), the electrodes montage consisted in placing the anode over the area F3 of the EEG 10-20 system and the return (cathode) electrode over the right supra-orbital area as reported in Figure 1c. The duration of the stimulation consisted of about 17 min (1040 s) with 15 s of currents fade-in and fade-out. Configuration of the sham condition included 15 s of fade-in, 10 s of actual current delivery and 15 s of fade-out given at the beginning of the experiment only (see Figure 1d).

### 2.5 Statistical analyses

As expected, the nature of our oddball task caused ceiling effects in the correct responses for all conditions (accuracy rate > 98%). All the trials that included either a false alarm or a missed response were left out from subsequent analyses on RT, as well as trials corresponding to RT faster than 150 ms or exceeding 1.96 standard deviations from the mean (number rejected trials: *M* = 3.14, *SD* = 1.39). All valid RT were then log-transformed to the base *e* in order to ensure a normal distribution of the data. We considered only trials having no missing values at the two main outcomes RT and PD, resulting in 52 trials overall. Importantly, these data points were not collapsed across conditions; hence *Trial* was included in the analyses as an independent fixed factor, and thus affording a greater reliability and robustness of the findings.

In order to study the effect of tDCS on the behavioral and physiological responses, we performed two linear mixed models (LMM) on RT and on PD as dependent variables. Individual (subject-specific) variation was accounted for by considering *Subjects* as random effect. Fixed effects, repeated within subjects, were specified for *Condition* (2 levels, *real* and *sham), Time* (2 levels, *T1* and *T2*) and *Trial* (52 levels); whereas *Order* (2 subgroups, *sham-real* and *real-sham)* was considered as a between-subject fixed effect. In addition, the interaction *Condition* x *Time* was assessed. Post hoc comparisons were adjusted with Sidak correction for multiple comparisons.

The above LMM were subsequently adjusted for subjective arousal (measured by STAI-Y State score) and for physiological arousal (evaluated by PrePD) in order to assess their effects on tDCS-induced modulation. Akaike information criteria (AIC) was used to select the best fitted models (the lower AIC the better model) and the corresponding predictors.

Finally, to control for any interdependence between the subjective and physiological measures of arousal, we calculated Pearson’s (r) two tailed correlations between PrePD and STAI-Y State score. Correlations coefficients were all non-significant (*p*’s > 0.05). All statistical analyses were conducted on SPSS Statistics (IBM Corp, Armonk, NY, USA).

## 3. Results

Despite the random assignment, the participants included in the two *Order* subgroups exhibited different levels of physiological tonic arousal (PrePD) already at the baseline of the first experimental session, that is before applying the tDCS electrodes (two tailed independent t-tests [t = −3.64, df = 11.82, *p* = .003]). No difference was found between the subgroups in the STAI-Y scores [t = −.41, df = 11.57, *p* = .68].

As for the reported sensations, a Wilcoxon matched pair test revealed no significant difference between sham and real stimulation [*Z* = 1.34, *p* = 0.18]. It was also ensured that their written responses were consistent with their oral report. Therefore, it was safe to assume that participants were completely unaware of the type of stimulation protocol.

### 3.1 Reaction times – RT

The unadjusted linear mixed model on RT [AIC = −2574] revealed no significant effects of the *Order* [*F_(1,11)_* = .59, *p* = .477] and a trend toward significance for *Condition* [*F_(1,2419)_* = 3.76, *p* = .053] and *Trial* [*F_(51,98)_* = 1.47, *p* = .05], with slower RT occurring at the end of each tasks. A significant effect of *Time* [*F_(1, 2399)_* = 12.15, *p* < .001] showed that performance significantly improved from *T1* [*M* = 5.98; *SE* = .05] to *T2* sessions [*M* = 5.96; *SE* = .05], indicating an overall practice effect. Importantly, we found a significant *Condition* x *Time* interaction effect [*F_(1,2403)_* = 12.08, *p* = .001], indicating a different trend for real and sham conditions. The post-hoc comparison for *Time* revealed a significant performance improvement during *sham* (*p* < .001), but not during *real* stimulation (*p* = .99). This finding suggests that real tDCS hindered the practice effect that was present in the sham condition.

Next, LMM adjusted for STAI-Y and PrePD were separately performed (see Supplementary Table 1). We found an overall significant contribution of STAI-Y [*F_(1,2312)_* = 9.44, *p* = .002] and more importantly a significant 3-way interaction [*Condition* x *Time* x STAI-Y: *F* = 19.1, df = 3/1857, *p* < .001], indicating that the subjective level of arousal affected the interaction *Condition* x *Time* on RT. Specifically, STAI-Y state scores were predictive of the performance variations across tDCS conditions. During sham session a performance improvement was observed for all the continuum of arousal, although it diminished as the level of STAY-Y increased. In the real tDCS condition, RT proved to be faster only when the levels of arousal were low, whereas such pattern was abolished or even reversed with higher levels of arousal (i.e., higher STAI-Y scores, see Figure 2).

**Fig. 2.**
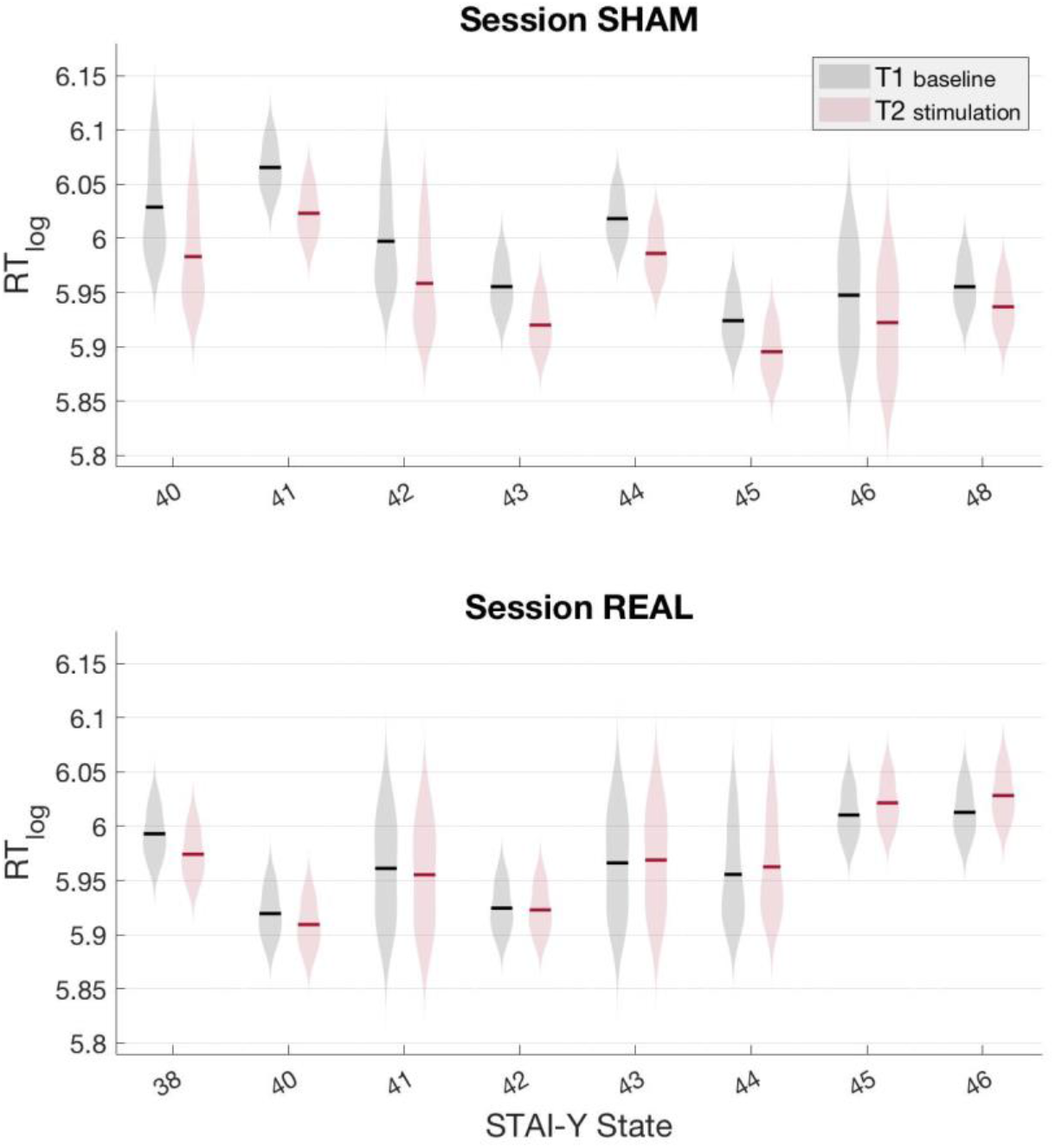
Reaction times by subjective arousal. Average log-based RT of model fitted values are plotted as a function of STAI-Y scores, with results from session sham (top panel) and real (bottom panel). Each mean value is marked over the corresponding distribution of the data. Colors grey and red represent the baseline (T1) and stimulation (T2) task, respectively.

After adjusting for PrePD, *Condition* and *Time* remained significant [*F_(1,2131)_* = 6.88, *p* = .009; *F_(1,1967)_* = 9.44, *p* = .032 respectively], although the physiological predictor did not reach statistical significance [PrePD: *F_(1, 1834)_* = 3.52, *p* = .061]. Also in this case, the 3-way interaction [*Condition* x *Time* x PrePD: *F_(3,1307)_* = 5.57, *p* = .001] revealed that the interaction between *Condition* and *Time* was affected by participants’ physiological level of arousal. Consistently with the aforementioned effects of subjective levels of arousal, RT improvement across time was consistent in the sham condition, but larger during trials with a reduced PrePD. During real tDCS, a trend toward lower or no improvement was observed as physiological arousal increased (see Figure 3).

**Fig. 3.**
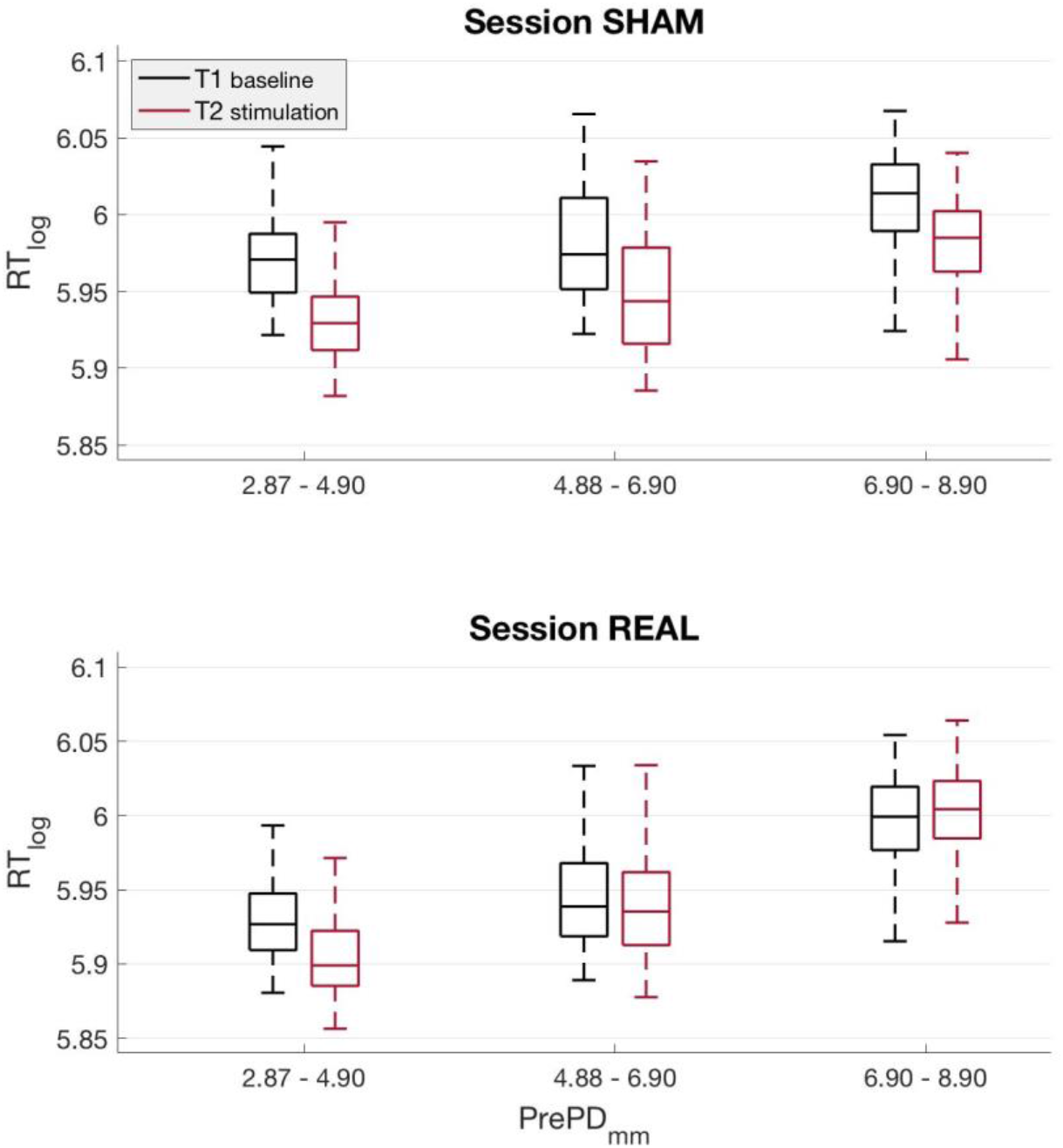
Reaction times by physiological arousal. On each box, the interquartile range, the whiskers and the median of predicted log-based RT are represented for three linearly interspaced bins of pre-target pupil diameter, with results from session sham (top panel) and real (bottom panel). Colors grey and red represent the baseline (T1) and stimulation (T2) task, respectively.

Based on the present results, a far more consistent trend emerged from the adjusted models as compared to the same raw data (see Figure 4). This finding corroborates the importance of not disregarding discrepancies rooted in interindividual differences, such as in physiological and subjective arousal, but rather include them as predictors along with individual random effects.

**Fig. 4.**
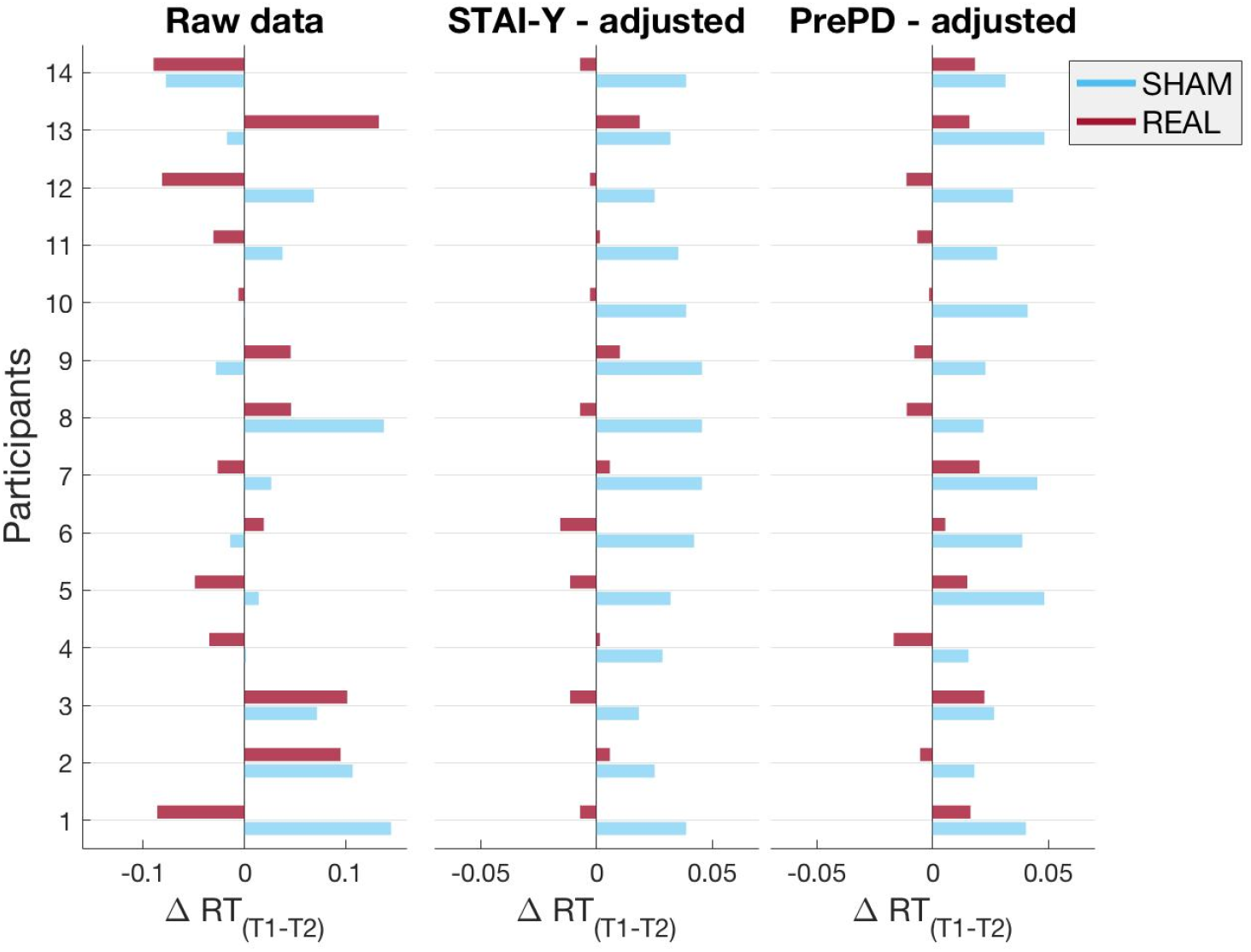
Subject variability of reaction time change. Average log-based RT differences between the baseline (T1) and stimulation (T2) tasks are plotted on the vertical axis for each participant, separately for session sham (blue bars) and real (red bars). A different bar plot is used to represent mean differences from raw (left panel) and fitted data from the adjusted models using STAI-Y (middle panel) and PrePD (right panel) predictors. Negative and positive values on the horizontal axis indicate slower and faster performance, respectively.

### 3.2 Pupil dilation – PD

In the unadjusted LMM on PD, [AIC = 964.66], all fixed effects were significant [*Condition: F_(1,1340)_* = 10.46, *p* = .001; *Time: F_(1,1445)_* = 15.83, *p* < .001; *Trial: F_(51,88)_* = 5.94, *p* < .001] except for the factor *Order* [*F_(1,11)_* = 1.96, *p* = .18] and the interaction between *Condition* and *Time* [*F_(1,1362)_* = .75, *p* = .38]. Importantly, pupil dilation decreased from *T1* [*M* = .375; *SE* = .023] to *T2* sessions [*M* = .34; *SE* = .023], indicating a general habituation of the phasic pupillary responses. However, no specific effect of tDCS on PD was revealed.

The adjustment for STAI-Y got worse the model fitting [AIC = 986.38], making the interaction *Condition* x *Time* x STAI-Y not significant [*F_(1,1066)_* = 1.08, *p* = .35] (see Supplementary Table 2). On the contrary, adjusting for PrePD strongly improved the model fitting [AIC = −365.48], with significant PrePD [*F_(1,1794)_* = 2231.23, *p* < .001] and interaction *Condition* x *Time* x PrePD effects [*F_(3,1172)_* = 6.5, *p* < .001]. In detail, during the sham condition a decrease in pupil dilation consistently occurred throughout the range of PrePD values, whereas during real tDCS the pupil dilation progressively shifted toward a maximal suppression during trials with larger PrePD (Figure 5).

**Fig. 5.**
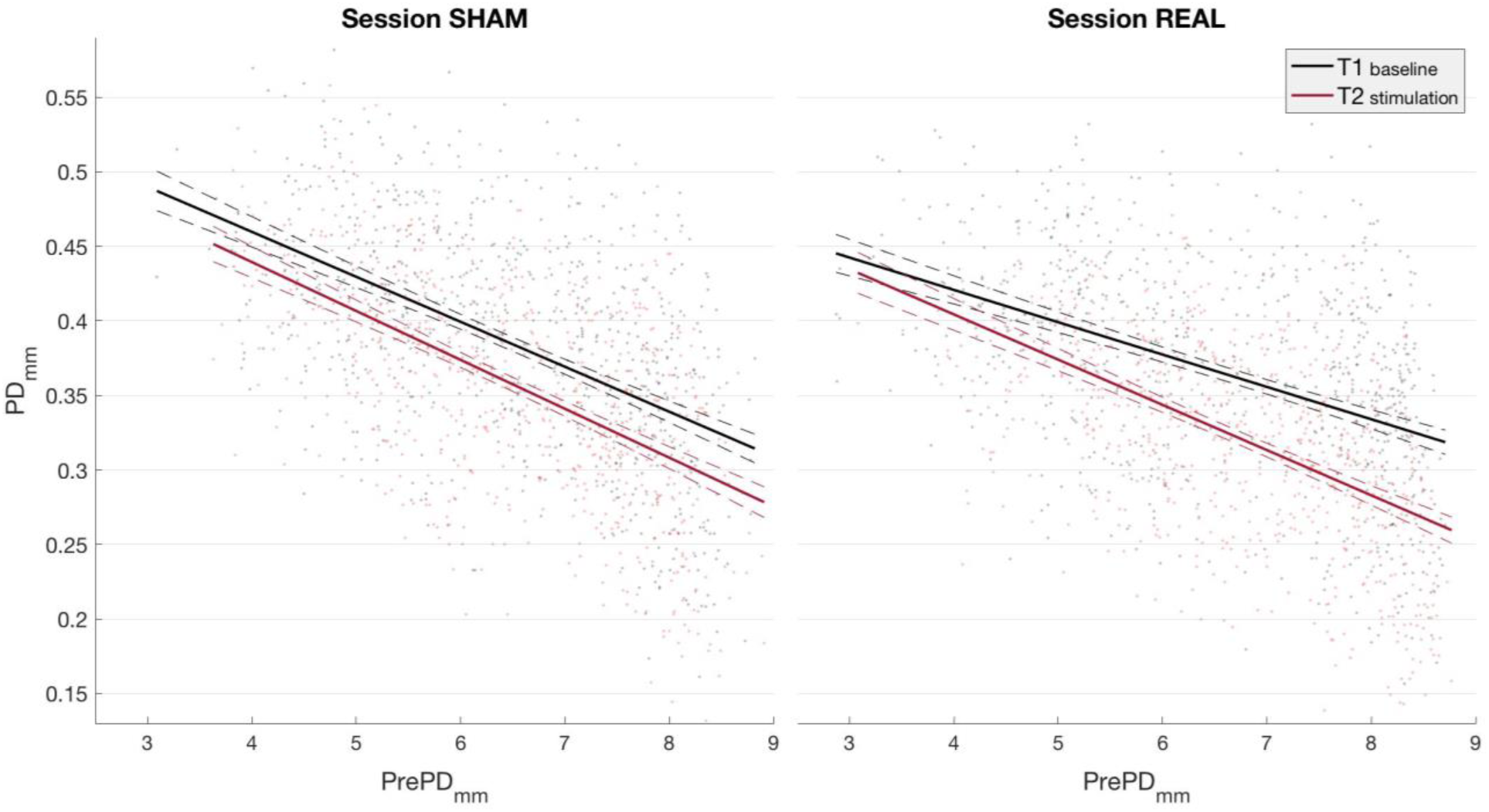
Pupil dilations explained by physiological arousal. Model fitted PD values are plotted against pre-target pupil diameter, with results from session sham (**left panel**) and real (**right panel**). Grey and red best-fitting lines describe the trend of pupil dilation data points over pre-target pupil diameter respectively for the baseline (T1) and stimulation (T2) task. Dashed lines represent prediction functional bounds, i.e. the uncertainty of predicting the fitted lines.

## 4. Discussion

In the present study, we addressed the question of whether variable effects of single session tDCS could be dependent on the degree of arousal experienced before and during the experiment.

Subjective and physiological levels of arousal significantly accounted for the variation of reaction times across two experimental sessions. Real tDCS appeared to hinder the practice effect observed during the sham condition, with a trend becoming especially evident at higher levels of arousal. As for pupil dilation, its values were significantly tied to the corresponding physiological fluctuations of arousal. In particular, a more reduced pupillary response emerged during real tDCS as arousal levels increased.

These results shed light on one relevant factor, which seems to account for the paucity of consistency across tDCS effects in some experiments. What effectively emerges is that arousal is predictive of the modulations induced by tDCS on task performance. A number of studies, which reported a considerable inter- and intra-individual variability in response to tDCS protocols, investigated the impact of demographic characteristics (e.g., age, gender), cortical architecture variations or physiological measures specific to the targeted areas (e.g., levels of excitability of the primary motor cortex), yet without considering general measures of activation comparable to arousal (9,64–68).

Here, we collected ratings on the subjective level of anxiety (i.e., STAI-Y State) before each experimental session, thus serving as a fixed measure of arousal. Pre-target pupil diameter was instead used as a dynamic proxy of arousal, allowing us to track its ongoing fluctuations (50,52,69). We confirmed that pupil dilation values were negatively related with pre-target pupil diameter across all conditions, as frequently reported in the literature (50,70–72).

When our measures of arousal were accounted for by statistical analyses, a clear picture emerged, indicating that the effects induced by tDCS on the behavioral responses were dependent on both subjective and physiological levels of arousal levels. In the sham session, participants speeded up their responses when they completed the task for the second time. This practice effect emerged somewhat independently of both the subjective and physiological levels of arousal, although a slightly more pronounced improvement appeared with lower levels in either measures. During the application of real tDCS, however, performance ceased to improve with the exception of trials characterized by smaller pre-target pupil diameter and participants with a lower score at the STAI-Y questionnaire. A negative or null behavioral outcome of anodal tDCS is not uncommon in the literature and learning impairments have been reported in a host of different tDCS studies involving specific learning outcomes, such as unimproved working memory for recognition or implicit categorization, blocked consolidation of visual perception and inhibited motor learning (68,73–78).

We chose response speed as behavioural measure, given that its intrinsic low sensitivity heavily relies on prior levels of fatigue and general activation (79–82). The interpretation of our behavioural results is arguably consistent with an inverted U-shape curve between task performance and arousal. According to this relationship, performance decline would occur when arousal levels are either too high or too low (33,83). None of the participants reported sleep deprivation or otherwise drowsiness-related conditions. Therefore, we can assume that the lower values of our predictors effectively corresponded to moderate and not low levels of arousal. With this in mind, the finding that facilitatory effects are principally associated with a moderate level of cortical excitation seems to support the proposed cellular mechanism for a cortical excitation-inhibition balance (16,84). On these grounds, tDCS exogenous modulation would negatively impact on the normal cortical functioning whenever the levels of endogenous neural activity increase to the extent of a dysfunctional neuronal gain, with spontaneous task disengagement causing slower responses. A direct consequence of this mechanism would be the inhibition of task learning effects, unless the endogenous system is sufficiently inactive, as in low arousal trials. The latter scenario provides an additional argument for when single session tDCS is found to improve task performance in the face of variable but otherwise moderate and well-balanced arousal levels. The understanding that an unbalanced combination of endogenous and exogenous excitability-increase events can, in fact, lead to negative effects is also coherent with frameworks on brain activity-dependent plasticity and on signal-to-noise ratio mechanisms (20,78). Results on pupil dilation, which represents a physiological response to relevant stimuli, corroborate the above interpretation. Only when the ongoing levels of arousal were considered in the analyses, a specific effect of tDCS on pupil dilatation was revealed. An overall reduction of pupil dilation occurred when participants completed the task for the second time (T2), consistently with a physiological habituation effect that paralleled the practice effect seen in the behavioral results (85,86). In particular, pupil dilation evenly decreased for the entire range of arousal in the sham session, but crucial variations emerged during the application of real tDCS: looking at the lower end of the arousal range, pupil dilation values were not as much reduced as in sham session. Conversely, a more pronounced reduction in pupil dilation was observed in trials associated with higher arousal. These, in fact, corresponded to the trials of unimproved response times following real tDCS. Therefore, habituation of a phasic response may not necessarily indicate the same outcome direction as the better performance after a practice effect (85,87). Pupil dilations primarily reflect the timely increase of neural gain control, which translates into a system’s responsivity amplification, and as such can be ascribed in the aforementioned inverted-U curve (34,50,71,88). The implication is that the additive effect of an exogenous neuromodulation would, on the one hand, contrast the natural habituation effect on pupil dilation occurring below the intermediate range of tonic arousal and, on the other hand, accentuate task disengagement at higher levels of tonic arousal, hence a greater reduction in phasic response. An analogous explanation was put forward in a recent tDCS work showing a reduction of pupil dilation - but no behavioral effects - during a Go-NoGo task, whereby it was argued that an offline tDCS enhancement of neuronal membrane potential could hinder or replace the endogenous gain control mechanisms of locus coeruleus (40).

Furthermore, outside the tDCS literature, phasic pupillary responses were found to be reduced whenever participants’ attention was not directed to the task, such as during episodes of mind wandering (72,89–91). Indeed, recent empirical and theoretical formulations of mind wandering have proposed that the locus coeruleus–norepinephrine system is tightly linked to different internally-driven cognitive states, i.e., on- and off-task states with various degrees of deliberate control (36,92). In this respect, the possibility of a direct and focally targeted tDCS modulation of mind wandering has been recently debated with uncertain conclusions (93). Based on this knowledge, it is not unlikely that our tDCS effects would also be partly dependent on the arousal-mediated propensity of mind wandering activity during the task. Although not covered by the aims of this study, the above possibility justifies the argument for a selective alteration of arousal via exogenous neuromodulation. For example, vigilance decrements and physiological sleep pressure were somewhat diminished after prolonged frontal anodal tDCS (41,94) with a magnitude of effects greater than caffeine (95,96). Whereas for the arousal modulation related to a specific event, stimulus-locked bursts of electric random noise stimulation were used to enhance performance and LC phasic responses, as indexed by skin conductance measurements (25). Despite this compelling evidence, it is still difficult to conclude that certain tDCS montages can directly modulate the deep brain arousal structures (97–99). Note though that other mechanisms could involve changes in the neocortical neurons, whose membrane potential shifts are known to be coupled with alteration in pupil diameter (34,100,101).

In summary, our data collectively offer an explanation for the negative or null effects of a common prefrontal tDCS application. We are aware that the interpretation of these particular results may not apply to all tDCS studies. Nevertheless, the large variability of arousal levels that we found across participants leads us to reflect more closely on what may mask the desired effects in the varied and still growing landscape of stimulation studies, which often fail to incorporate, but simply acknowledge, the crucial aspect of individual state-dependent variables (102–105).

The importance of brain state is not a novel idea in the literature on non-invasive brain stimulation. The ongoing or basal levels of activation, included in the concept of “state-dependency”, have been extensively reported to impact the effects of transcranial magnetic stimulation (TMS) (106). Nevertheless, considering the mechanisms of action of tDCS, which modulates excitability of neurons by hyperpolarizing or depolarizing their membrane potential (107,108), tDCS effects might be more sensitive to the arousal levels than TMS. In a similar vein, these considerations might be applicable to any kind of current stimulation modality.

Taken together, the discussed findings should encourage a more careful interpretation of null or negative effects of tDCS. This is far from saying that all replication failures are due to inherently inefficacious tDCS protocols with exhausted future potential (15). Such observations should instead help ascribing the outcome of those protocols to the interrelation of the locus coeruleus–norepinephrine system and the spreading of induced currents in the brain (40,42). In this sense, future tDCS studies might consider useful to have both dynamic and fixed measures of arousal as an accurate way to monitor its impact on the final outcome. If successful, these achievements would be of great help also in assessing the degree of effectiveness with which tDCS protocols are being utilized to treat or ameliorate clinical conditions.

## Acknowledgements

CF, CM and DB have been supported by the projects of the Italian Ministry of Health “Ricerca Corrente”.

