## Supplementary Material for "Arousal levels explain inter-subject variability of neuromodulation effects"

### Control experiment

A control experiment was conducted to exclude the eventuality that our tDCS protocol could produce any subjective discomfort, thus generating different alerted states across tDCS conditions. These preliminary results served as validation of the stimulation protocol to which participants were to remain blind and without any confounded pupillary dynamics.

*Participants*

Twelve right-handed healthy participants took part in this control study (mean age = 26.38; *SD* = 1.85; 9 females). Data of two subjects were rejected prior to analyses due to the excessive noise in her/his pupil signal (i.e. interpolation rate > 30% of the whole epoch). Note, though, that the reduced sample size did not affect the counterbalancing of either gender or stimulation order.

Participants had no history of neurological or psychiatric illness and had normal or corrected-to-normal visual acuity. Ethical approval was obtained by the Ethics Committee of the IRCCS Centro San Giovanni di Dio Fatebenefratelli, Brescia, Italy. All participants were given written informed consent.

*Experimental procedure*

All participants underwent real and sham tDCS in the same day, interposed by a pause of 15 minutes. Each protocol of stimulation lasted about 5 minutes, a duration whose excitability modulations have been proved to return to baseline within the following 5 minutes (1,2). The stimulation condition order was randomly assigned and counterbalanced across participants. The testing session consisted in recording pupillary dynamics at rest using the exact same setting as in our main study. After each session, participants were given a questionnaire to rate their perceived sensations or discomforts (3,4).

Pupil signal acquisition and preprocessing were conducted with the same set up and by following the same steps as in the main study. Eye blink correction was implemented with a custom script in MATLAB (MathWorks, USA). A shape-preserving piecewise cubic interpolation method was chosen to interpolate values ranging from 70 ms before blink onset to 300 after blink offset.

Exactly the same stimulation parameters as in the main study were used (current intensity = 1mA; electrodes = 35 $cm^2$; montage = F3 - right supra-orbital area), except for the duration of the stimulation, which was 390 s in the real tDCS (including 10 s of fade-in and fade-out) and in the sham tDCS (10 s of fade-in, 10 s of actual current delivery and 10 s of fade-out, both before and following 330 s of null current).

*Results and discussion*

Firstly, the rate of interpolated data (i.e. full or half-eye blinks), which is known to tap into the dopaminergic and fatigue-related neural pathways, was used to measure participants' discomfort (5–7). No significant difference was found between the interpolation rates within sham (8.07% ± 5.8) and anodal tDCS condition (7.76% ± 5.9) [$t_{(9)} = .47$; $p = 0.64$].

Secondly, we run Wilcoxon matched pair tests to analyze responses to the questionnaire, yielding non-significant differences between stimulation protocols in any of the probed sensations (all $p > 0.1$). These results were also consistent with participants' oral report, which confirmed the relatively weak, hence comparable degree of discomfort during both conditions.

### Main experiment

**Behavioral performance (RT)**

| **Models** | **AIC** | **Fixed Factors** | $\mathbf{F_{(df)}}$ | ***p* value** |
|---|---|---|---|---|
| 1) Unadjusted model | -2574 | *Trial* | $1.47_{(51,98)}$ | 0.050 |
| | | *Time* $^{*}$ | $12.15_{(1,2399)}$ | <0.001 |
| | | *Order* | $0.59_{(1,11)}$ | 0.477 |
| | | *Condition* | $3.76_{(1,2419)}$ | 0.053 |
| | | *Condition* x *Time* $^{*}$ | $12.08_{(1,2403)}$ | <0.001 |
| 2) Adjusted for STAI-Y | -2588 | STAI-Y $^{*}$ | $9.44_{(1,2312)}$ | 0.002 |
| | | *Trial* $^{*}$ | $1.56_{(51,99)}$ | 0.030 |
| | | *Time* $^{*}$ | $4.08_{(1,2284)}$ | 0.043 |
| | | *Order* | $0.59_{(1,11)}$ | 0.454 |
| | | *Condition* $^{*}$ | $41.82_{(1,2318)}$ | <0.001 |
| | | *Condition* x *Time* x STAI-Y $^{*}$ | $19.10_{(3,1857)}$ | <0.001 |
| 3) Adjusted for PrePD | -2549 | PrePD | $3.52_{(1,1834)}$ | 0.061 |
| | | *Trial* $^{*}$ | $1.54_{(51,98)}$ | 0.031 |
| | | *Time* $^{*}$ | $4.62_{(1,1967)}$ | 0.032 |
| | | *Order* | $0.39_{(1,12)}$ | 0.542 |
| | | *Condition* $^{*}$ | $6.88_{(1,2131)}$ | 0.009 |
| | | *Condition* x *Time* x PrePD $^{*}$ | $5.57_{(3,1307)}$ | 0.001 |

Table 1.

The table illustrates the structure and statistics of the models utilized for the dependent variable reaction time. All models include subjects as random factor. Significant fixed factors are displayed with an asterisk.

**Pupil dilatation (PD)**

| **Models** | **AIC** | **Fixed Factors** | **$F_{(df)}$** | ***p* value** |
|---|---|---|---|---|
| 1) Unadjusted model | 965 | *Trial* * | $5.94_{(51,88)}$ | <0.001 |
| | | *Time* * | $15.83_{(1,1445)}$ | <0.001 |
| | | *Order* | $1.96_{(1,11)}$ | 0.180 |
| | | *Condition* * | $10.46_{(1,1340)}$ | 0.001 |
| | | *Condition* x *Time* | $0.75_{(1,1362)}$ | 0.380 |
| 2) Adjusted for STAI-Y | 986 | STAI-Y * | $11.98_{(1,1248)}$ | 0.001 |
| | | *Trial* * | $6.03_{(51,89)}$ | <0.001 |
| | | *Time* | $0.19_{(1,1321)}$ | 0.666 |
| | | *Order* | $2.61_{(1,11)}$ | 0.133 |
| | | *Condition* | $2.15_{(1,1276)}$ | 0.142 |
| | | *Condition* x *Time* x STAI-Y | $1.08_{(3,1066)}$ | 0.350 |
| 3) Adjusted for Pre-PD | -365 | PrePD * | $2231.23_{(1,1794)}$ | <0.001 |
| | | *Trial* * | $2.84_{(51,87)}$ | <0.001 |
| | | *Time* * | $8.21_{(1,1888)}$ | 0.004 |
| | | *Order* * | $11.47_{(1,12)}$ | 0.005 |
| | | *Condition* * | $7.85_{(1,2158)}$ | 0.005 |
| | | *Condition* x *Time* x PrePD * | $6.51_{(3,1172)}$ | <0.001 |

Table 2.

The table illustrates the structure and statistics of the models utilized for the dependent variable pupil dilation. All models include subjects as random factor. Significant fixed factors are displayed with an asterisk.
